# SCEMENT: Scalable and Memory Efficient Integration of Large-scale Single Cell RNA-sequencing Data

**DOI:** 10.1101/2024.06.27.601027

**Authors:** Sriram P Chockalingam, Maneesha Aluru, Srinivas Aluru

## Abstract

**Motivation:** Integrative analysis of large-scale single cell data collected from diverse cell populations promises an improved understanding of complex biological systems. While several algorithms have been developed for single cell RNA-sequencing data integration, many lack scalability to handle large numbers of datasets and/or millions of cells due to their memory and run time requirements. The few tools which can handle large data do so by reducing the computational burden through strategies such as subsampling of the data or selecting a reference dataset, to improve computational efficiency and scalability. Such shortcuts however hamper accuracy of downstream analyses, especially those requiring quantitative gene expression information.

**Results:** We present SCEMENT, a SCalablE and Memory-Efficient iNTegration method to overcome these limitations. Our new parallel algorithm builds upon and extends the linear regression model previously applied in ComBat, to an unsupervised sparse matrix setting to enable accurate integration of diverse and large collections of single cell RNA-sequencing data. Using tens to hundreds of real single cell RNA-seq datasets, we show that SCEMENT outperforms ComBat as well as FastIntegration and Scanorama in runtime (upto 214X faster) and memory usage (upto 17.5X less). It not only performs batch correction and integration of millions of cells in under 25 minutes, but also facilitates discovery of new rare cell-types and more robust reconstruction of gene regulatory networks with full quantitative gene expression information.

**Availability and implementation:** Source code freely available for download at https://github.com/AluruLab/scement, implemented in C++ and supported on Linux.

**Supplementary information:** Supplementary data are available at *Bioinformatics* online.

## 1 Introduction

Several different methodologies have been developed for integrating multiple single cell RNA-sequencing (scRNA-seq) datasets, with the aim of eliminating batch effects inherent in samples spanning different locations, labs, and conditions, while also conserving biological variation. Currently available scRNA-seq integration methods can be classified into three major categories: (i) Methods that output embedding onto a reduced dimensional space such as PCA (Xu *et al*., 2021; Korsunsky *et al*., 2019), (ii) Methods that output graphs such as a cell-cell k-nearest-neighbor graph (Polański *et al*., 2020; Haghverdi *et al*., 2018), and (iii) Methods that retain the gene-level quantitative information, i.e., their output is a gene expression matrix containing gene expression profiles from input cells (Zhang *et al*., 2019; Johnson *et al*., 2007). Although these methods have been useful in integrating single cell datasets generated under a variety of tissues and conditions, their applicability is constrained by limitations on scale of data they could handle, and thus integration of large numbers of cells and complex scRNA-seq datasets still remains a challenge.

A recent comprehensive survey of 16 different supervised and unsupervised scRNA-seq integration methods showed that none of these methods could successfully integrate 970K cells collected from multiple samples of a mouse brain dataset due to runtime and memory constraints (Luecken *et al*., 2022). To improve computational efficiency and scalability, some tools designed for large data integration perform one or more of the following: (i) partition the data at discrete steps of the processing pipeline to solve data-specific problems (Li *et al*., 2022), (ii) operate on a reduced dimensional space of cells (Korsunsky *et al*., 2019; Haghverdi *et al*., 2018), (iii) use only a representative subset of datasets/genes (such as a reference dataset or a few highly variable genes) (Dhapola *et al*., 2022; Hao *et al*., 2023), and (iv) use unscaled data instead of scaled data to avoid generating a dense matrix (Luecken *et al*., 2022). However, such shortcuts result in sampling bias and limit applicability of the integrated data for downstream processing steps, especially those that require a complete gene expression matrix such as for gene regulatory networks and cell-cell interaction networks.

Our goal is to overcome limitations on numbers of genes or cells, while simultaneously achieving data size scalability and adequate performance. Here, we present a novel approach that uses a sparse implementation of the empirical Bayes-based linear regression model to integrate scRNA-seq data from a large number of datasets and expression profiles. While the concept of applying linear regression model is well established in various biological research areas (Pierson and Yau (2015), Kerr (2003), Dirmeier *et al*. (2018)), including in single cell research (Johnson *et al*., 2007)), our method SCEMENT (SCalablE and Memory-Efficient iNTegration) incorporates multiple algorithmic improvements for a faster and efficient way to enable large-scale scRNA-seq data integration with millions of cells and tens of thousands of genes.

In this paper, we demonstrate that by engineering sparsity during all computations, even in cases where mathematical expressions involving the input sparse matrix *X* could potentially lead to intermediate dense matrices, and by designing an efficient order of computations, SCEMENT outperforms ComBat (Johnson *et al*., 2007), FastIntegration (Li *et al*., 2022), and Scanorama (Hie *et al*., 2019) in run-time (upto 214X faster) and memory usage (upto 17.5X less). It performs batch correction and integration of 4 million cells collected from 121 samples with more than 38K genes in just 22 minutes. In addition, SCEMENT not only maintains meaningful biological gene expression variations across cell-types even when cells are clustered by their condition, but also facilitates downstream processing of single cell data for better identification of rare cell-types and more robust reconstruction of gene networks with full gene expression information.

## 2 Methods

Linear regression models provide two key advantages for integration of gene expression profiles from scRNA-seq data: 1) t he a bility to accommodate various experimental conditions and parameters, and 2) the ability to retain quantitative gene expression values after integration. To account for technical and biological variations resulting from different conditions, a linear model uses numerical and categorical type variables to represent these variations. In this work, we assume that a given dataset includes only categorical variables as is common with single-cell RNA-seq datasets.

### 2.1 SCEMENT’s approach for large-scale integration

Similar to the model outlined in ComBat (Johnson *et al*., 2007), we start with the following generative empirical Bayes-based linear regression model:

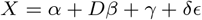

where the gene expression data *X* of *m* cells and *n* genes is modeled as a linear function of four terms:

- The average or overall gene expression, denoted by *α*, is an *m × n* matrix. Each row *i* in *α* corresponds to the gene expression profile of the set of conditions the cell *i* belongs to.
- Linear combination of the independent variables, *β*. It is the matrix of regression coefficients (size *c × n*) with each column corresponding to a specific variable. In case of integration, the independent variables are the condition/batch variables. *D* is the design matrix, a binary *m × c* matrix such that entry *D*[*i, j*] is 1 if input *i* is observed under the condition *j*.
- Additive batch effect denoted by *γ*.
- Multiplicative batch effect denoted by *δ*. Furthermore, *ϵ* is the error term, assumed to follow a normal distribution with variance *σ*, i.e, *ϵ* ∼ 𝒩 (0, *σ*^2^*I*_*m*_).

SCEMENT employs an efficient algorithm to estimate the model parameters: *α, β, γ, δ* and *σ*. Complete details of the empirical Bayes solution are provided in the supplementary text. In this section, we present the key contributions underlying SCEMENT that enable large-scale integration. As is common in statistics, we use 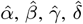 and 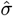 to represent the estimated values of *α, β, γ, δ* and *σ*.

#### 2.1.1 SCEMENT’s strategy for computing the standardized matrix

Integration of gene expression data starts with the computation of the standardized matrix *Z* (see Supplementary text S1.1.1 for how *Z* is defined and computed) from the input data matrix *X*, since *Z* provides well-behaved mean and variance characteristics for more robust analyses. Computing *Z* requires computation of 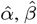 and the variance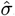, and these require matrix computation operations involving the data matrix and the design matrix. For microarray and bulk RNA-seq datasets, where the number of observations are in the order of thousands, currently available dense matrix multiplication routines are sufficient. However, for single-cell RNA-seq datasets with hundreds of thousands to millions of cells, computation of *Z* necessitates a memory-efficient approach.

##### Computing 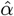 and 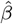

We propose a space and time efficient method to accelerate the computation of 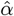, with the following optimization strategies that enable scaling to millions of cells and tens of thousands of genes.

1. Current implementations use a 32-bit integer matrix to represent the design matrix *D*. When all the *β* variables are only categorical, entries of *D* are binary *i*.*e*., either 1 or 0 and hence do not require 32-bit representation. We use 8-bits for entries in *D* to improve space by a factor of 4, which also leads to time efficient computation of *D*^*T*^ *D*.
2. For computing 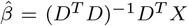, the order of multiplying the matrices can have significant effect on the runtime. For real datasets, since the number of observations is an order of magnitude larger than the number of genes, multiplication of (*D*^*T*^ *D*)^−1^ with *D*^*T*^ *X*^*T*^ is a better way to compute 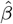 when compared to the product of (*D*^*T*^ *D*)^−1^*D*^*T*^ and *X*^*T*^ because the former takes *O*(*c*^2^) time while the latter takes *O*(*c*^2^*m*) time when *m* ≫ *n*.

Subsequent to computing 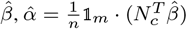 is computed by using a simple matrix vector multiplication routine.

##### Computing 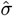

Computing 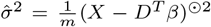 · 𝟙_*m*_ can lead to explosion in the amount of memory usage even when *X* is sparse, because realizing *X* − *D*^*T*^ *β* in-memory creates a dense intermediate matrix of size *m × n*.

Let *R* = *D*^*T*^ *β*. To avoid realizing this intermediate matrix *X* − *R*, SCEMENT employs the algebraic expansion (*X* − *R*)^⊙2^ · 𝟙_*m*_ = *X*^⊙2^ · 𝟙_*m*_ + *R*^⊙2^ · 𝟙_*m*_ − (2*X* ⊙*R*) · 𝟙_*m*_ (similar to (*a*−*b*)^2^ = *a*^2^ + *b*^2^ − 2*ab*). By expanding (*X* − *R*)^⊙2^ into three terms, SCEMENT computes the three terms efficiently as follows.

1. The first term *X*^⊙2^ · 𝟙_*m*_ is computed by squaring each entry in sparse *X* summing it row-wise. Since *X* is a sparse matrix, this can be computed using sparse matrix routines.
2. To compute the second term, we exploit the unique property of the matrix *R* that *R* has as many unique rows as the number of unique condition-profiles. A condition-profile is the set of unique conditions a cell can belong to. Even in a large collection of datasets comprising millions of cells, the number of unique condition-profiles is limited to few dozens, guaranteeing sparsity. For efficient computation, we exploit the fact that all the cells that belong to the same condition-profile have the same row vector in *D* and, therefore in *R*. In order to compute *R*^2^ · 𝟙_*m*_ efficiently, SCEMENT first enumerates all the unique condition-profiles *D* and then computes the product vector 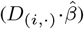 separately for each one of them. We accomplish this with the aid of two auxillary data structures: (a) A configuration matrix *G*, which contains set of condition-profile vectors, and (b) A condition-profile lookup vector *l*_*D*_. *R*^⊙2^ · 𝟙_*m*_ can be computed by adding these vectors as many times as the number of cells in the corresponding condition-profiles.
3. The condition-profile lookup vector *l*_*D*_ and the configuration matrix *G* can also be used to compute the third term efficiently. SCEMENT employs a series of multiplication of sparse-matrix and dense-vector, one for each unique condition-profile.

With our efficient way of computing these three terms, the partial sums corresponding to individual condition-profiles are added as soon as they are computed, thus saving memory and time.

Finally, in order to maintain low memory footprint, we do not fully realize *Z* as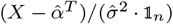. We retain the left-hand side 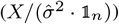 and the right-hand side 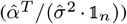 of *Z* as matrices *X*^*′*^ and *M*^*′*^ respectively.

Each of the above steps to compute (*X* − *R*)^⊙2^ can be accomplished efficiently in parallel as follows. For the first term *X*^⊙2^ · 1_*m*_, each of the row sum of squares can be computed by a parallel reduction across each each row *X*^⊙2^. In case of the second term, a coordinate sparse (COO) representation of the matrix *X* along with the row-wise distribution of configuration matrix *G* and the look-up vector *l*_*D*_, enables efficient distribution of the row-wise computations in parallel. Similarly parallel computation of the third term is accomplished by row-wise distribution of the configuration matrix *G*.

Note that *X*^*′*^ is the same size as the input matrix and *M*^*′*^ is of the size *c*^*′*^ *× n*, where *c*^*′*^ is the number of unique configuration-profiles in the dataset. In other words, *M*^*′*^ has only one row only for each unique configuration-profile and not for each cell. In the next sections, we show how *X*^*′*^ and *M*^*′*^ matrices can be used in the downstream computations instead of the *Z* matrix.

#### 2.1.2 SCEMENT’s approach for Batch Correction

##### Iterative 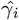 and 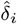 update

Empirical Bayes method follows an iterative algorithm to estimate 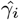 and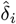. The primary challenge in the iterative update of 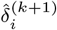 at the *k*-th iteration is to evaluate the expression

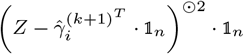

without realizing the dense matrix *Z* in memory, where 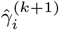 is the update of *γ*_*i*_ at *k*-th interation.

Similar to the computation of 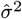 discussed in section 2.1.1, the above computation can be accomplished by expanding the expression into three terms (i) *Z*^⊙2^ · 𝟙, 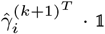, and (iii)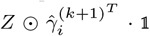. As mentioned earlier, in SCEMENT, *Z* is maintained as two matrices *X*^*′*^ and *M*^*′*^. Therefore, the first and the third terms expand to (*X*^*′*^ − *M*^*′*^)⊙2 · 𝟙 and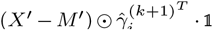. Both these terms can be further expanded and each of the individual terms can be computed without having to realize the standardized *Z* matrix. Each term is successively added up to obtain the update for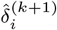.

By not directly computing the *Z* matrix and using algebraic expansion for the terms where *Z* appears, we retain sparsity of the computations, and thereby efficiently compute each update. Also, parallel computation of these terms is accomplished in a similar manner to the computations described in section 2.1.1.

##### Batch Corrected Matrix

In the final step, we convert the sparse *X* matrix to dense and update it as the batch corrected matrix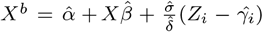. Similar to computation of *Z* in section 2.1.1, it is possible to retain the batch corrected matrix, *X*^*b*^ as two sparse matrices – one each corresponding to the left-hand side and the right-hand side of *Z*. However, to facilitate downstream processing of the integrated matrix such as for computing PCA, UMAP, t-SNE, clustering and plotting, SCEMENT converts the sparse *X* to dense *X*.

### 2.2 SCEMENT’s Implementation

We implemented two versions of SCEMENT compatible with the *AnnData* data structure used in *Scanpy*. One in the *python* programming language (pySCEMENT) and the second is a faster parallel version in *C++* (SCEMENT−CPP). Both versions use single precision floating point (32-bit) values for the computations. The *python* version uses sparse matrix libraries available in the *scipy* python package for representing the input data and *X*^*′*^. Though the input data and *X*^*′*^ are stored in compressed sparse array (CSR) representation, *numpy* library arrays are used to store all the other matrices and vectors in the algorithm. For the construction of design matrix *D*, we used the *formulaic* library, which allows for saving space with the use of 8-bit integers.

OpenMP is used for implementation of the parallel SCEMENT algorithm in *C++*. In order to enable efficient computations, we use the coordinate sparse (COO) representation to store the input data and *X*^*′*^. *Armadillo* C++ libraries are used for representing all other dense vectors and matrices. While parallel sparse computations are implemented as per section 2.1, *ScaLAPACK* library is used for computations involving dense matrices and vectors.

### 2.3 Performance Assessment of SCEMENT

We performed two types of evaluation studies using real scRNA-seq datasets from different tissues and organisms (Table S1). First, we compared performance of four other previously published integration methods–FastMNN, ComBat, Scanorama and Seurat (Table S1) in terms of both integration quality and separation of clusters in UMAP plots using scRNA-seq datasets from *A. thaliana* plant root (Jean-Baptiste *et al*., 2019; Gala *et al*., 2021) and human aortic valve (Xu *et al*., 2020). Results from these runs were subsequently used to compute quality control metrics according to the **scIB** software package (Luecken *et al*., 2022). We also visually compared clusters of aortic valve dataset generated using SCEMENT, with the aforementioned four methods. Cell-type annotations used in UMAP plots of human aortic valve cells were according to Xu *et al*. (2020), and for Arabidopsis cells according to Jean-Baptiste *et al*. (2019), and Gala *et al*. (2021).

Next, we evaluated scalability of the integration methods using two different human peripheral blood mononuclear cell (PBMC) datasets: 1) a COVID-19 dataset of about 1.23 million cells from 205 samples (Ren *et al*., 2021) and, 2) a collection of 17 human PBMC datasets containing a total of 794,170 cells obtained from the 10X Genomics web repository (Table S2). We assessed runtime and memory usage of all the integration methods with varying number of datasets/cells, and with integrated data consisting of union as well as intersection of genes. All runs were conducted on a machine equipped with a 72-core Intel^®^ Xeon^®^ E7-8870 CPU and main memory of 1 TB shared between all the cores.

#### 2.3.1 Construction and analysis of Gene Regulatory Networks

We used the pySCENIC workflow (Kumar *et al*., 2021) to construct gene regulatory networks (GRNs) from integrated data matrices consisting of union as well as intersection of genes. Here, we use human PBMC datasets (Table S2) containing cells ranging from ≈ 20,000 to 166,000 cells. The quality and performance of the resulting networks were assessed using standard statistical measures: recall (percentage of correct edges predicted), precision (percentage of correct edges among all edges inferred), and the F-score defined as: F-score = (2 *×* precision *×* recall)*/*(precision +recall). To evaluate biological relevance of networks generated from different integrated matrices, we used the known human transcriptional regulatory network from the TRRUSTv2 database (Han *et al*., 2018) as ground truths. This network was constructed by text-mining of published literature and manual curation, and include a total of 8,426 high confidence regulatory interactions between 795 transcription factors (TFs) and 2492 target genes (Table S3). For the purpose of computing statistical measures, all known TF-target interactions from amongst the 8426 interactions were considered as true positives (TPs) whereas TF-target interactions not listed in the TRRUSTv2 database were considered as true negatives (TNs).

#### 2.3.2 Cell-type Identification from Large-scale Integrated Data

We generated integrated data matrices with SCEMENT-CPP using varying number of datasets and cells sampled from the ≈ 1.2 million cell scRNA-seq dataset (Ren *et al*., 2021), and subsequently applied the Azimuth package (https://azimuth.hubmapconsortium.org/) on the integrated data to automatically identify various human PBMC cell-type populations.

## 3 Results and Discussion

### 3.1 Linear regression model for scRNA-seq integration

SCEMENT is designed to be an unsupervised computational method that makes large-scale batch correction and integration of scRNA-seq datasets feasible, while retaining gene expression profiles of all available genes from input cells in the integrated data matrix. A recent survey of over 16 different supervised and unsupervised scRNA-seq integration methods (Luecken *et al*., 2022) ranked four unsupervised methods − FastMNN (Haghverdi *et al*., 2018), Seurat v3 (Butler *et al*., 2018), Scanorama (Hie *et al*., 2019) and ComBat (Johnson *et al*., 2007), amongst the top 10 best performing methods. These four methods also meet our criteria of returning an integrated gene expression matrix of batch corrected values as output. Therefore, we sought to further evaluate FastMNN, Scanorama, Suerat and ComBat.

We used scRNA-seq datasets generated from two different organisms containing varying sizes and complexity to assess the four integration methods: wild-type *A. thaliana* plant root dataset (AtRD) with 14,427 cells collected from two separate studies and nine different batches (Jean-Baptiste *et al*., 2019; Gala *et al*., 2021), and a human aortic valve dataset (HAVD) containing 17,985 cells collected from four individuals; two healthy and two diseased (Xu *et al*., 2020).

We employed 8 different evaluation metrics from the scIB package (Luecken *et al*., 2022) in conjunction with UMAP visualizations to make valid comparisons of the four integration methods. Our results show that Seurat outperforms the other three methods in executing batch correction and integration with respect to HAVD dataset, while ComBat performs better overall with the AtRD datasets (Table 1). Our studies further show that the ComBat model preserves biological variation within different cell type/states more strongly than the other three methods (Figure 1; Figure S1). Whereas cell-types such as lymphocytes, macrophages, endothelial cells (VEC) and interstitial cells (VIC) are separated into well-defined clusters by all four methods, UMAP visualizations show that Scanorama, Seurat and FastMNN have a tendency to overmix cells, thus resulting in poor representation of the transcriptional heterogeneity between healthy and diseased cell states. In this context, it should be noted that the Arabidopsis dataset consists of cells from only wild-type root samples (normal) whereas the aortic valve dataset consists of cells from healthy (normal) and diseased (abnormal) individuals. We therefore speculate that the sub-optimal performance of ComBat with the aortic valve dataset as observed in Table 1 is perhaps due to the less aggressive cell mixing characteristics of the ComBat model, which in turn facilitates a better separation of healthy from diseased cells, even within the same cell-type. Moreover, ComBat requires significantly less runtime and memory to integrate these scRNA-seq datasets when compared to Seurat (Table 2). Based on these results, we decided to use the linear regression model as a basis for developing a faster and more efficient method for large-scale scRNA-seq integration.

**Table 1:** Benchmarking ComBat, Scanorama, Seurat and FastMNN for scRNA-seq data integration using eight scIB metrics (Luecken *et al*. (2022)). A brief description of the metrics is given in Supplementary text. Numbers in bold represent the average of the 8 metrics for each method.

|  | Aortic Valve Dataset |  |  |  | Arabidopsis Dataset |  |  |  |
| --- | --- | --- | --- | --- | --- | --- | --- | --- |
|  | ComBat | Scanorama | Seurat | FastMNN | ComBat | Scanorama | Seurat | FastMNN |
| NMI_cluster/label | 0.3694 | 0.3569 | 0.4807 | 0.3931 | 0.7384 | 0.7323 | 0.6981 | 0.5833 |
| ARI_cluster/label | 0.2013 | 0.2293 | 0.3753 | 0.3418 | 0.6299 | 0.6345 | 0.5655 | 0.4035 |
| ASW_label | 0.4974 | 0.5033 | 0.5327 | 0.5004 | 0.5568 | 0.5547 | 0.5542 | 0.5171 |
| ASW_label/batch | 0.8499 | 0.8892 | 0.9101 | 0.8907 | 0.9198 | 0.9044 | 0.8926 | 0.9083 |
| Isolated F1 | 0.5191 | 0.4940 | 0.7298 | 0.4495 | 0.7496 | 0.8534 | 0.7470 | 0.7568 |
| Isolated ASW | 0.3942 | 0.4428 | 0.5640 | 0.4451 | 0.6278 | 0.6055 | 0.6297 | 0.5489 |
| Graph Conn. | 0.9404 | 0.9605 | 0.9882 | 0.9057 | 0.9486 | 0.9607 | 0.9613 | 0.9727 |
| HVG Cons. | 0.1340 | 0.0205 | 0.0340 | 0.0490 | 0.2697 | 0.0163 | 0.0654 | 0.0203 |
| **Average Score** | **0.4882** | **0.4871** | **0.5769** | **0.4969** | **0.6801** | **0.6577** | **0.6392** | **0.5889** |

**Table 2:** Runtime and memory usage of ComBat, Scanorama, Seurat and FastMNN for scRNA-seq data integration. HAVD = Human aortic valve dataset and AtRD = Arabidopsis root dataset.

|  | Runtime (s) |  | Memory (GB) |  |
| --- | --- | --- | --- | --- |
|  | HAVD | AtRD | HAVD | AtRD |
| ComBat | 172.81 | 66.71 | 19.41 | 6.11 |
| Scanorama | 530.75 | 125.63 | 38.34 | 13.24 |
| Seurat | 3298.44 | 1792.36 | 84.53 | 42.63 |
| FastMNN | 187.22 | 136.12 | 5.01 | 4.66 |

**Fig. 1:**
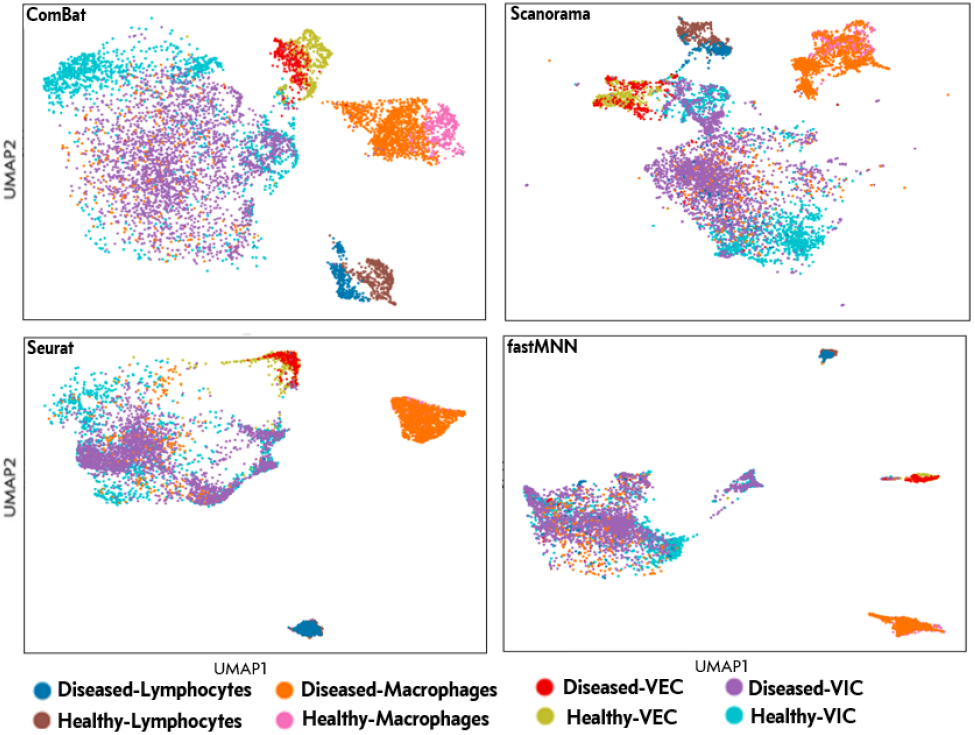
UMAP visualizations of cell-type clusters from the human aortic valve dataset after batch correction and integration with ComBat, Scanorama, Seurat and FastMNN

### 3.2 Overview of the SCEMENT algorithm

We developed SCEMENT by making the following algorithmic improvements to the empirical Bayes-based linear regression model: (i) a sparse implementation of the Empirical Bayes-based integration method, (ii) maintaining sparsity of matrices throughout and avoiding dense intermediate matrices through algebraic manipulation of the matrix equations, (iii) employing an efficient order of operations that allows for accelerated computation of the batch integrated matrix, and (iv) a scalable parallel implementation that enables integration of diverse datasets to more than four millions cells.

The overall approach incorporated in the SCEMENT algorithm is illustrated in Figure 2. Our method takes as input single cell data matrices along with the batch and other condition labels (for example, gender or the disease condition of the patients) for all experiments. However, in contrast to other integration methods, SCEMENT provides an optional pre-processing step (shown as Step P of Fig. 2). This step allows for construction of an integrated data matrix containing the union of genes across all batches. For each of the sample/batch, we first identify genes missing in that particular sample/batch, but present in any of the other batches. We then insert rows with zero entries corresponding to the missing genes into each of the gene expression matrices such that all of the input matrices have the same set of genes. Subsequent merging of the modified matrices, thus generates an integrated data matrix containing the union of genes.

**Fig. 2:**
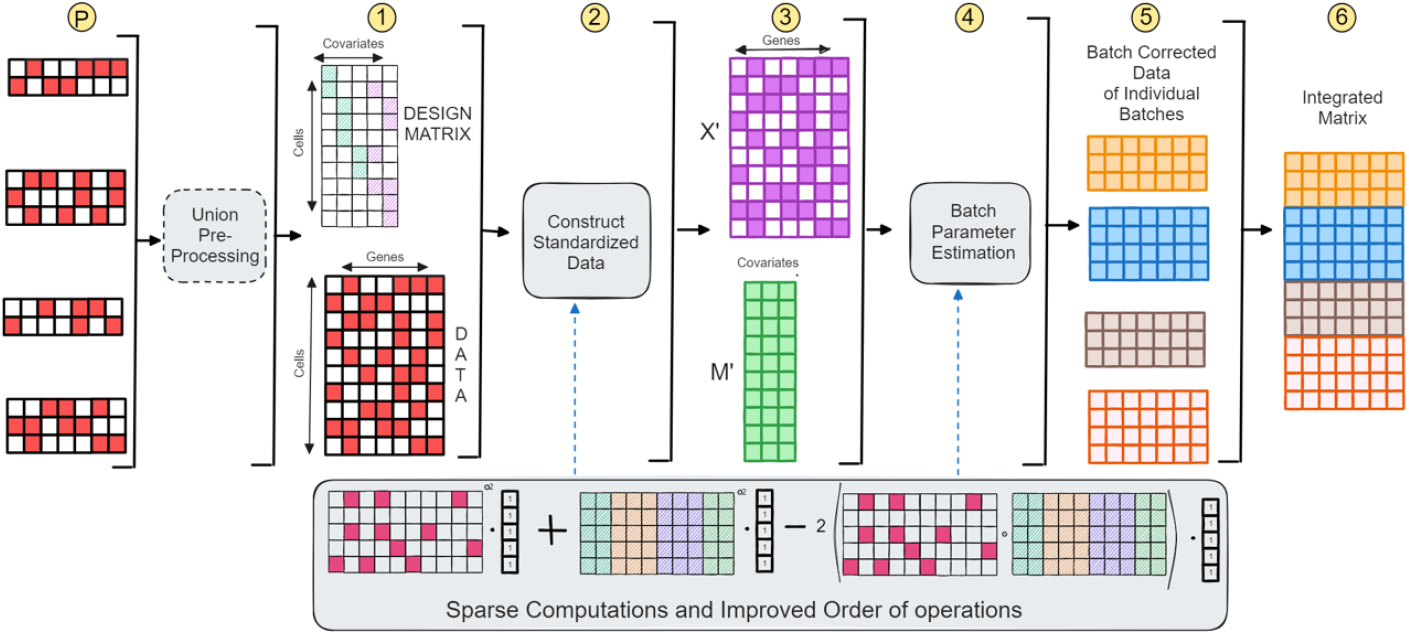
Overall worfklow of SCEMENT. A detailed description of the integration methodology is provided in sections 2.1 and 3.2

We next compute the standardized matrix *Z* by exploiting the limited number of configurations present in the design matrix, in comparison to the millions of cells present in the large data. In SCEMENT, *Z* is computed as a combination of the matrix pair (*X*^*′*^, *M*^*′*^): the average expression matrix and the pooled variance per configuration (Step 3 of Fig. 2). The use of the matrix pair aids in faster computation of the gene-level and condition-level parameters of the underlying regression model. Subsequently, each batch is processed separately to estimate batch effects. Estimation of batch effects also relies on the (*X*^*′*^, *M*^*′*^) representation. After adjusting each individual matrix for batch effects, the final matrix is generated in the last step as an integrated cell X gene matrix of all the batch corrected matrices. These algorithmic improvements in SCEMENT achieve speed and improved scalability without any detrimental effect on the quality of scRNA-seq integration when compared to ComBat (Figure S2).

### 3.3 SCEMENT enables large-scale scRNA-seq integration

We assessed SCEMENT’s performance for large-scale integration of scRNA-seq datasets by measuring its runtime and memory usage with varying number of datasets/batches and cells. We used two sets of PBMC derived scRNA-seq datasets – 205 samples from COVID-19 patients (Table 3) and another from 17 different healthy individuals (Table S2). We performed batch correction and integration using intersection (genes common to all datasets) as well as the union of genes (all genes from all datasets), for all of the 8 subsets ranging from 3 to 205 COVID-19 datasets, and 7 subsets ranging from 2 to 17 healthy PBMC datasets. As expected, Table 3 and Table S2 show that as the number of datasets increase, the number of intersecting genes decrease. In contrast, this number increases when we use the union of genes for constructing the integrated data.

**Table 3:**
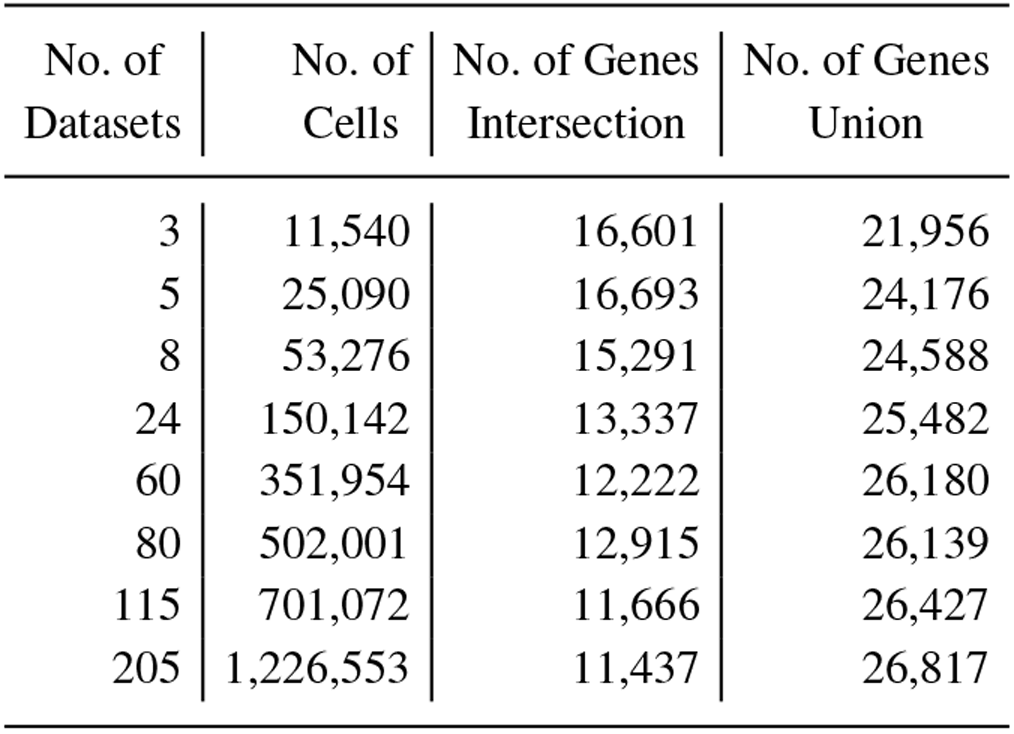
scRNA–seq datasets from Ren *et al*. (2021)

| No. of Datasets | No. of Cells | No. of Genes Intersection | No. of Genes Union |
| --- | --- | --- | --- |
| 3 | 11,540 | 16,601 | 21,956 |
| 5 | 25,090 | 16,693 | 24,176 |
| 8 | 53,276 | 15,291 | 24,588 |
| 24 | 150,142 | 13,337 | 25,482 |
| 60 | 351,954 | 12,222 | 26,180 |
| 80 | 502,001 | 12,915 | 26,139 |
| 115 | 701,072 | 11,666 | 26,427 |
| 205 | 1,226,553 | 11,437 | 26,817 |

We assessed runtime and memory consumption of both the python (pySCEMENT) and the C++ (SCEMENT-CPP) versions of SCEMENT and compared these with three other methods – ComBat, Scanorama and FastIntegration (Table S1). ComBat uses the linear regression model for scRNA-seq data integration (Johnson *et al*., 2007), and Scanorama (Hie *et al*., 2019) and FastIntegration (Li *et al*., 2022) have previously been shown to scale to million(s) of cells. FastIntegration is also a fast and high-capacity version of the Seurat integration tool. By default, currently available integration methods, including ComBat, Scanorama, and FastIntegration, generate an integrated data matrix containing cells with either a set of highly variable genes or the intersecting set of genes from all batches. However, in this study, we modified the ComBat workflow and applied the same preprocessing step as in the SCEMENT workflow (step P in Figure 2) to generate an integrated data matrix containing union of genes using ComBat. Therefore, we include ComBat, but exclude FastIntegration and Scanorama from our comparisons involving union of genes.

Our results show that SCEMENT-CPP outperforms all other methods in runtime and memory usage for both union as well as the intersection of genes (Figures 3 and 4; Tables S4–S7). It is upto 214X faster than FastIntegration, 184X faster than Scanorama, and 20X faster than ComBat depending on the number of datasets and genes involved in the integration task. Moreover, SCEMENT-CPP uses upto 12X less memory than Scanorama and 10X less than ComBat, thus enabling integration of more than a million cells and more than 26K genes in just 16-17 minutes. Even for smaller integration tasks for less than 200K cells where all other methods are able to complete the integration task, SCEMENT is about 10X times faster, uses less than 20GB of memory, and thus can accomplish this task on a modestly equipped workstation. Interestingly, ComBat and pySCEMENT perform comparably with respect to runtime, even though ComBat’s implementation is in parallel while pySCEMENT is sequential. pySCEMENT is also significantly more memory-efficient than ComBat. In fact, ComBat could not scale beyond 700K cells for the union of genes as it runs out of memory available on our benchmarking hardware. This is because pySCEMENT uses a sparse matrix with 32-bit floating point option while ComBat’s implementation uses the dense 64-bit matrix.

**Fig. 3:**
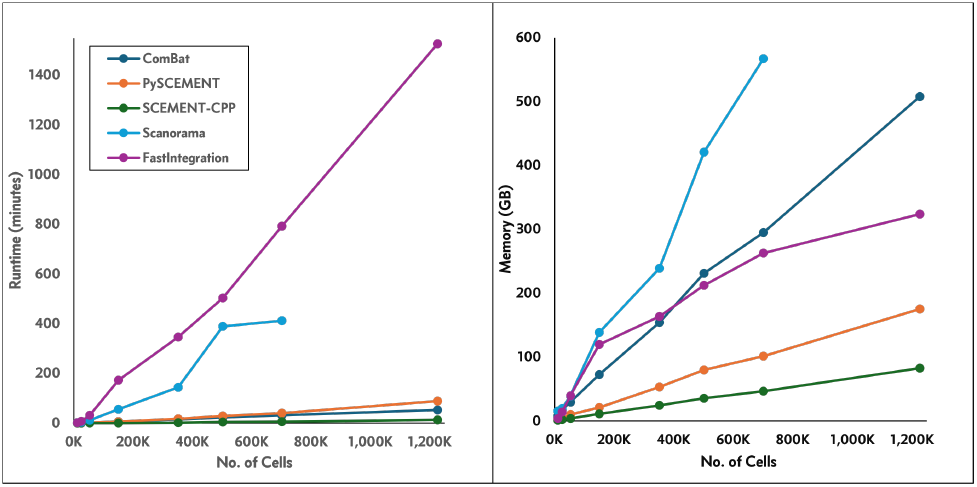
Runtime and memory usage of scRNA–seq data integration methods for intersection of genes.

**Fig. 4:**
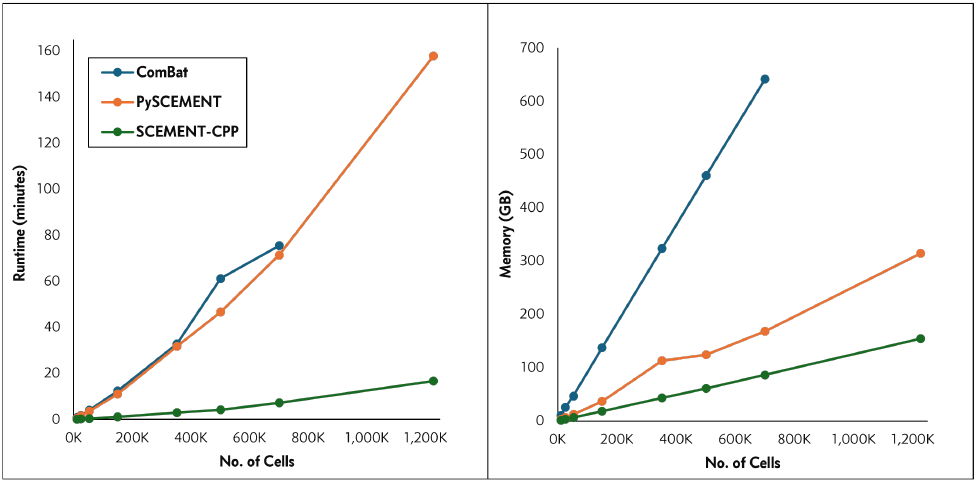
Runtime and memory usage of scRNA–seq data integration methods for union of genes.

FastIntegration and Scanorama require significantly longer runtimes and more memory usage respectively, when compared to SCEMENT and also ComBat (Figures 3 and 4; Tables S4–S7). In our studies, Scanorama could not successfully complete the runs beyond ≈800K cells for the intersection of genes. In addition, we (this study) and others have shown that FastIntegration can scale to millions of cells, however it accomplishes large-scale integration by 1) splitting a large integration task into a number of smaller integration tasks, which it then successively integrates to build the final integrated matrix, 2) restricting the data integration process to intersection of genes, and 3) requiring each individual dataset to be small as it uses Seurat to process the datasets. Of the 17 different human PBMC datasets (Table S2), one dataset contains a large number of cells (≈606,606 cells). In such a scenario, FastIntegration fails to complete the integration task.

To further assess SCEMENT’s scalability beyond a million cells, we applied it to a dataset of ≈ 4 million cells and 38,481 genes collected from 121 samples (Cao *et al*., 2020). Pre-processing and filtering of the data from 121 samples using *Scanpy* took about 78 minutes for pySCEMENT and SCEMENT-CPP, and while both were able to successfully integrate data from all samples, SCEMENT-CPP was signficantly faster than pySCEMENT and completed the run in just 22 minutes (Table S8).

### 3.4 SCEMENT enables identification of rare cell types from large-scale scRNA-seq data

Large-scale scRNA-seq data analysis has been shown to facilitat a deeper understanding of the cellular heterogeneity and discovery of new rare cell-types from complex tissues (Jindal *et al*., 2018; Qian *et al*., 2023). To assess whether SCEMENT enables improved identification of rare cell-types, we used scRNA-seq data from PBMCs (Ren *et al*., 2021), and generated integrated matrices from 3 random subsets with cells ranging from ≈ 50K to 1.2 million cells. The resulting matrices were subjected to automatic cell-type identification using the Azimuth package. It is currently not feasible to run Azimuth on large data with full gene set. Therefore, we restricted the number of genes in the integrated matrices to only the top 1000 highly variable genes to make it feasible to perform cell-type annotation on large-scale data. Even with such limitations, our results show that the number of cell-types identified increase with increasing number of cells (Table S9). Dendritic cells are the rarest cell-types amongst the PBMCs. In our study, a minimum of 500K cells were needed for discovery of cDC1 (conventional dendritic Cell 1) cells and more than a million for ASDC (AXL+dendritic cell) cells.

### 3.5 SCEMENT facilitates robust GRN reconstruction from integrated scRNA-seq data

Reconstruction of GRNs from high-throughput gene expression (for example, scRNA-seq) data requires quantitative gene expression information from a large number of genes and observations for determining accurate gene pair associations (Emmert-Streib *et al*., 2012). However, scRNA-seq data suffers from data sparsity, with each individual dataset containing gene expression profiles of a only a few thousand genes. Large-scale integration of multiple datasets may help overcome such limitations. It is currently not feasible to reconstruct GRNs from large-scale scRNA-seq data with hundreds of thousands to millions of cells and tens of thousands of genes, with existing GRN reconstruction methods. Therefore, to show utility of the integrated matrices and construct GRNs in a reasonable amount of time, we selected small-scale PBMC data with cells ranging from 20K to 166K (Table S2), and ≈ 2723 genes representing a non-redundant set of transcription factors (TFs) and target genes from the TRRUSTv2 manually curated gene regulatory network (Table S3).

We contructed GRNs from integrated data matrices containing the intersection (SCEMENT-CPP, ComBat, Scanorama, and FastIntegration) and the union set of genes (SCEMENT-CPP and ComBat) using the pySCENIC workflow.

Network quality evaluation measures (Table 4) show that SCEMENT and ComBat are comparable in their performance with respect to recall, precision and F-score values. Both these methods also outperform FastIntegration and Scanorama when using matrices containing intersecting set of genes. Furthermore, the higher ratio of TPs to total number of edges in these GRNs suggests less number of false positives in the network Table S10). More importantly, GRNs constructed from matrices containing the union set of genes perform significantly better than those constructed with matrices consisting only the intersecting set of genes. In fact, our results show that recall and precision increase with increasing number of cells and genes for networks containing the union of genes, while these measures decrease for networks reconstructed using the intersection of genes. (Figure 5 and Table S10). Overall, these results suggest that by incorporating gene expression profiles of all available genes from large number of input datasets in the integrated data matrix, SCEMENT enables more robust and accurate GRN reconstruction from single cell data.

**Table 4:**
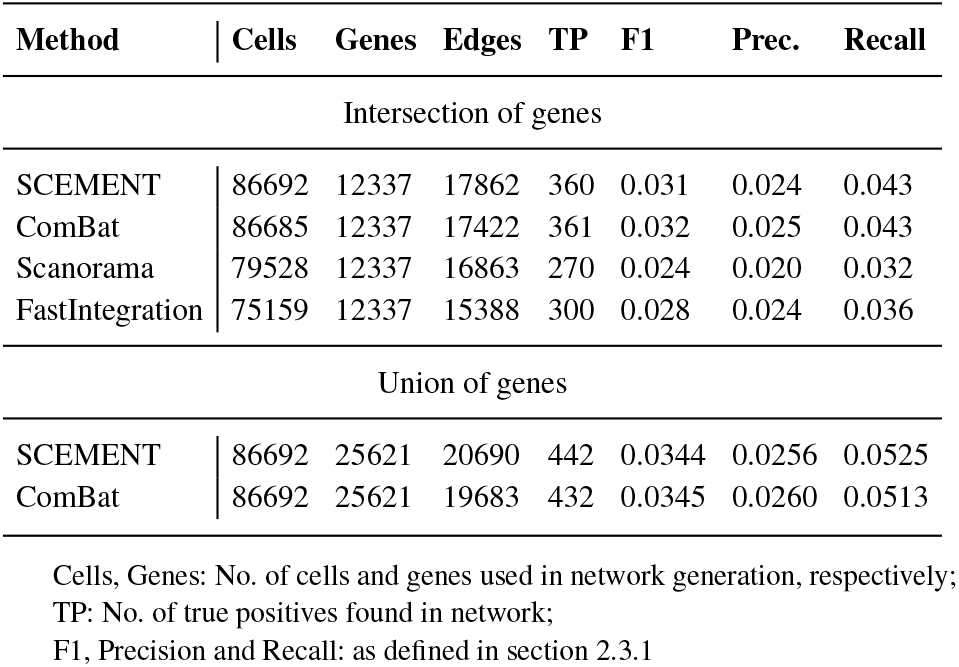
Assessment of network quality. scRNA-seq data from 9 different human PBMC datasets and cells totaling to ≈ 86K was used to generate six different integrated matrices containing either the intersection or the union of genes. GRNs were then reconstructed from each of these integrated matrices using the pySCENIC workflow.

**Fig. 5:**
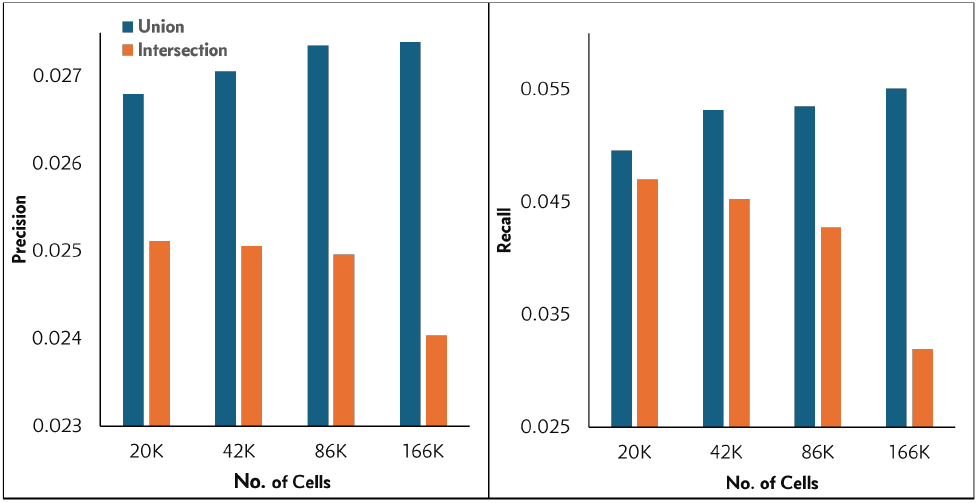
Precision and recall for networks generated from integrated matrices for union of genes with increasing number of cells.

## 4 Conclusions

Single cell transcriptome analyses are hampered by data sparsity, and large-scale integration of scRNA-seq data can overcome these limitations to provide a more comprehensive understanding of the cellular heterogeneity. We have developed a fast, scalable, and memory efficient method (SCEMENT) that enables accurate and large-scale integration of homogeneous and heterogeneous scRNA-seq datasets, and demonstrated its applicability on up to 4 million cells. SCEMENT is much faster and uses much less memory compared to existing methods. In fact, with SCEMENT, it is often not even necessary to have a high-memory system and an integration task of up to 500K cells and 25K genes can easily be completed on a laptop. We further demonstrate SCEMENT’s utility in the discovery of new and rare cell-types, and for more accurate and robust recontruction of large GRNs. Thus, SCEMENT is a simple but effective solution applicable to large and genome-scale integration of multiple scRNA-seq datasets, and opens new avenues for data-driven construction of atlas-scale cell maps.

## Acknowledgements

This work is supported in part by the National Science Foundation under NSF-2233887.

## Data Availability

The data used for evaluation of the proposed method are available in Zenodo, at *https://zenodo.org/doi/10.5281/zenodo.11521687*, and the accession numbers and the data sources are listed in Zenodo.

