## Supplementary Text S1 for "SCEMENT: Scalable and Memory Efficient Integration of Large-scale Single Cell RNA-sequencing Data"

### S1 Supplementary Text

#### S1.1 Empirical Bayes Model

The linear model for integration used in ComBat is represented by

$$X = \alpha + D\beta + \gamma + \delta\epsilon$$

where gene expression data is modeled as a function of the four terms.

- The average or overall gene expression, denoted by $\alpha$, is an $m \times n$ matrix. Each row $i$ in $\alpha$ corresponds to the gene expression profile of the set of conditions the cell $i$ belongs to.
- Linear combination of the independent variables, $\beta$. It is the matrix of regression coefficients (size $c \times n$) with each column corresponding to a specific variable. In case of integration, the independent variables are the condition/batch variables. $D$ is the design matrix, a binary $m \times c$ matrix such that entry $D[i, j]$ is 1 if input $i$ is observed under the condition $j$.
- Additive batch effect denoted by $\gamma$.
- Multiplicative batch effect denoted by $\delta$. Furthermore, $\epsilon$ is the error term, assumed to follow a normal distribution with variance $\sigma$, i.e, $\epsilon \sim \mathcal{N}(0, \sigma^2 I_m)$.

The Empirical Bayes based batch correction, as developed in COMBAT Johnson *et al.* (2007), for the above linear model proceeds with the following three steps:

1. Compute estimates $\hat{\beta}$, $\hat{\alpha}$ and standardize the input matrix.
2. Iteratively update to estimate batch effect $\delta$.
3. Compute the batch corrected matrix $X'$.

##### S1.1.1 Standardizing data

A standardized matrix is constructed first and used in all the subsequent steps since it provides well-behaved mean and variance characteristics for more robust downstream analyses.

A single sample-type can be observed simultaneously under multiple conditions. For example, a plant leaf sample belonging to a specific genotype can be observed under low and high stress conditions. Row $i$ in the $D$ matrix corresponds to the set of conditions under which sample $i$ is observed. We call this row as the condition-profile of sample $i$. In the case when a sample in $X$ belongs to exactly one condition, the least squares estimates for $\hat{\beta}$ and $\hat{\alpha}$ are

$$\hat{\beta} = (D^T D)^{-1} D^T X \quad \text{and} \quad \hat{\alpha} = \frac{1}{n} \mathbb{1}_m \cdot (N_c^T \hat{\beta})$$

respectively, where $N_c = [n_1, \ldots, n_c]^T$ is a vector with the number of observations in each condition, and $\mathbb{1}_m$ is $m$-length vector of ones. The estimate for the variance vector $\hat{\sigma}^2$ is

$$\hat{\sigma}^2 = \frac{1}{m}(X - D^T\hat{\beta})^{\odot 2} \cdot \mathbb{1}_m,$$

where the notation $A^{\odot 2}$ represents the Hadamard product of the matrix $A$ with itself, i.e., the entry $A^{\odot 2}[i, j]$ is $(A[i, j])^2$ and the dot product $A \cdot 1_m$ indicates the sum across the columns of $A$.

After estimating $\hat{\alpha}$ and $\hat{\sigma}$, the standardized matrix $Z$ is constructed as

$$Z = \frac{X - \hat{\alpha}^T}{\hat{\sigma} \cdot \mathbb{1}_n}$$

In the above equation, we use the division operation of two matrices of same sizes to indicate an element-wise division of numerator matrix entry to the denominator matrix entry.

In cases when a sample is subjected to multiple different conditions, let $S_c$ be the set of conditions that divides the dataset into $b$ partitions, i.e., $|S_c| = b$. In single-cell experiments, this can be the batch a sample belongs to. We assume that the first $b$ columns of $D$ correspond to these $b$ conditions. In this case, the estimates of $\hat{\beta}$ and $\hat{\sigma}^2$ remain the same as the first case, while $\hat{\alpha}$ is computed as follows:

$$\hat{\alpha} = \left(\frac{1}{n}\mathbb{1}_m \cdot (N_b^T \hat{\beta}_b)\right) + D_{-b}\hat{\beta},$$

where $\hat{\beta}_b$ is a $b \times m$ sub-matrix of $\hat{\beta}$ such that $\hat{\beta}_b = \hat{\beta}[(1, \ldots, b);\ (1, \ldots, n)]$ and $D_{-b}$ is $D$ with the entries corresponding to the batch rows set to 0, i.e., $D_{-b}[(1, \ldots, m); (1, \ldots, b)] = 0$.

##### S1.1.2 Iterative Algorithm for Batch update

Given that the dataset $X$ is merged from $b$ batches, then $Z$, in this case, is a standardized matrix that takes into account the batch weights. The linear model assumes that, for a given batch $i$ and gene $g$, the additive and multiplicative effects $\gamma_{ig}$ and $\delta_{ig}$ are drawn from $\mathcal{N}(X_i, t_i^2)$ and Inverse Gamma$(a_i, b_i)$ distributions respectively.

An empirical Bayes approach to estimate these parameters lends itself to an iterative solution. In iteration $k + 1$, the estimates for each batch $i$ are updated based on the estimates of the previous iteration $k$ ($\hat{\gamma_i}^{(k)}$ and $\hat{\delta_i}^{(k)}$) as follows Johnson *et al.* (2007). First, $\hat{\gamma_i}^{(k+1)}$ is computed as $\hat{\gamma_i}^{(k+1)} \leftarrow (t_i n \hat{\gamma_i}^{(k)})/(t_i n + \hat{\delta_i}^{(k)})$. Then, $\hat{\delta}_i$ is updated as

$$\hat{\delta}_i^{(k+1)} \leftarrow \frac{(2 + a - 1)}{2n}\left(b + \left(Z - \hat{\gamma}_i^{(k+1)^T} \cdot \mathbb{1}\right)^{\odot 2} \cdot \mathbb{1}\right),$$

where the dot product with $\mathbb{1}$ indicates the row-wise summing operation. The above steps are repeated until the percentage change of both $\hat{\gamma}$ and $\hat{\delta}$ is less than the required tolerance.

##### S1.1.3 Batch Corrected Matrix

After the $\alpha$, $\beta$, $\delta$, and $\sigma$ values are computed, the final step is to update the input gene expression matrix $X_b$.

$$X_b \leftarrow \hat{\alpha} + X\hat{\beta} + \frac{\hat{\sigma}}{\hat{\delta}}(Z_i - \hat{\gamma}_i)$$

#### S1.2 Metrics for Batch Integration

1. *NMI* and *ARI* compare the overlap of clustering with respect to the cell-type labels, with 0 being bad overlap and 1 being perfect match.
2. *ASW* measures the separation of clusters where 1 denotes dense and well-separated clusters, while 0 or -1 represents overlapping clusters.
3. *Isolated* scores were developed by Luecken *et al.* (2022) to evaluate how well data integration methods handle cell-types that appear in few batches.
4. *Graph Conn.* metric assesses how well the kNN graph constructed from integrated data directly connects all the cells of the same cell-type.
5. *HVG Cons.* score is a proxy for the preservation of highly variable genes after integration.
