## Supplementary figures and images for "SCEMENT: Scalable and Memory Efficient Integration of Large-scale Single Cell RNA-sequencing Data"

### Supplementary figure S1

#### Slide 1
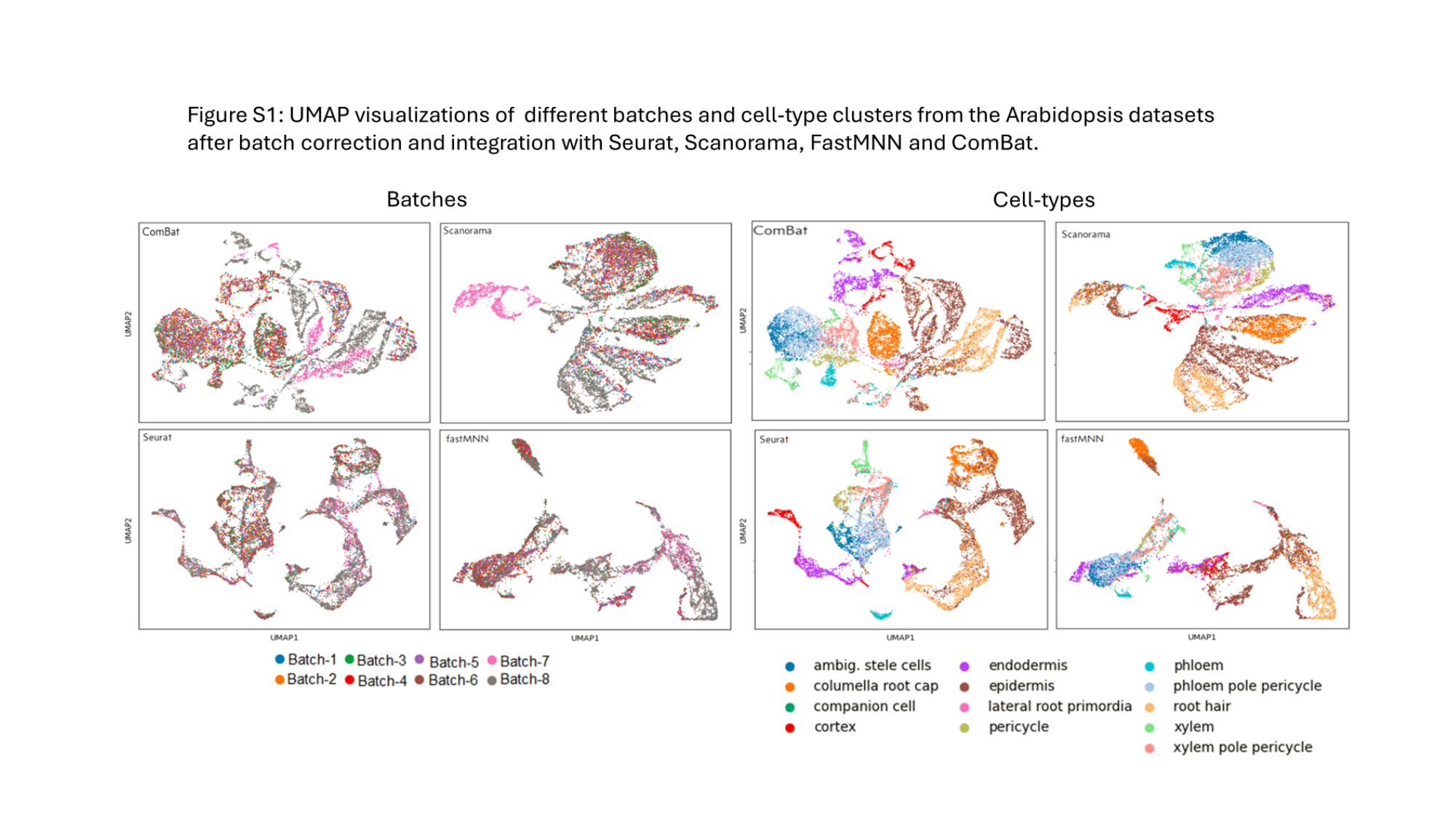

### Supplementary figure S2

#### Slide 1
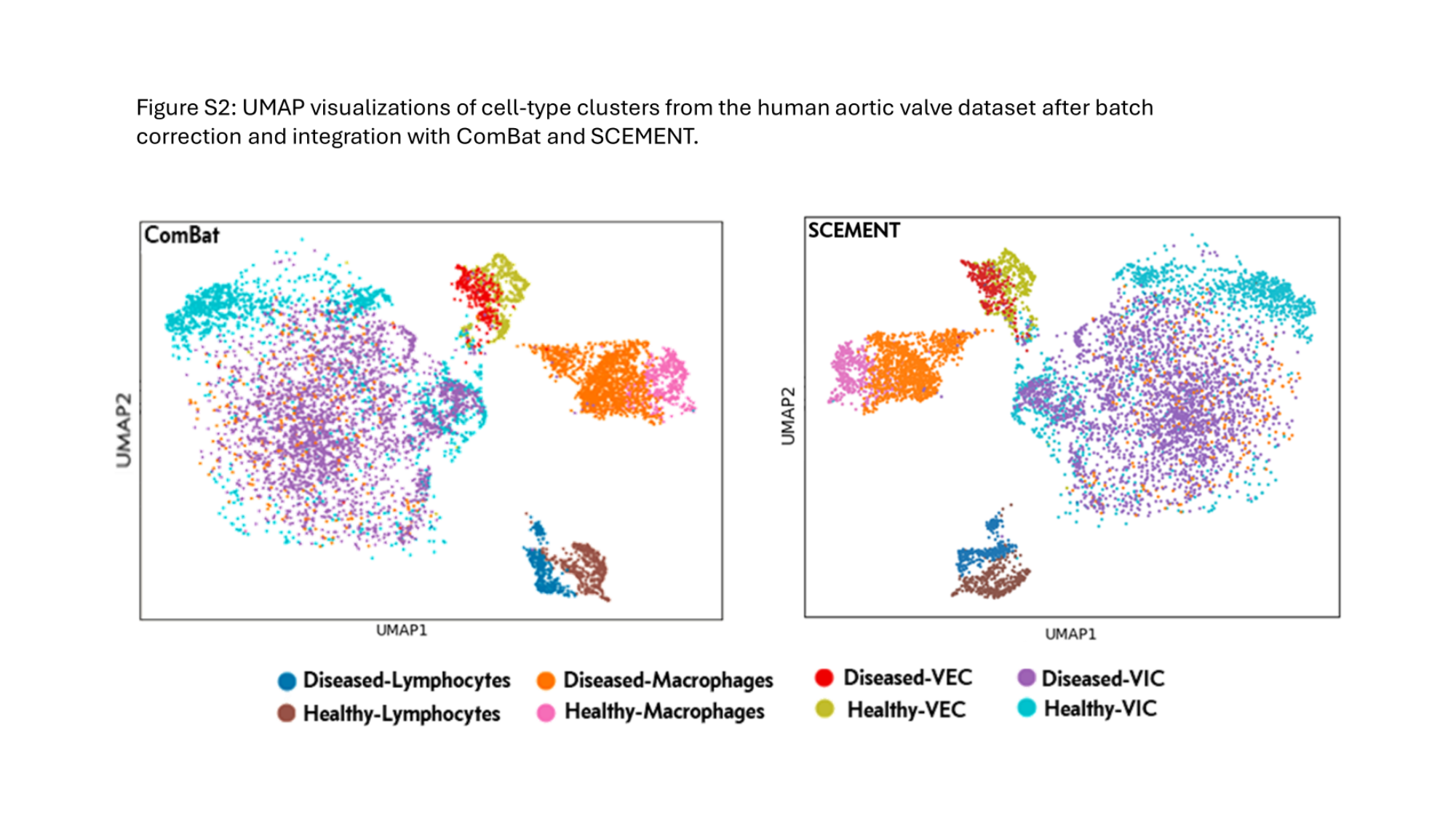

### Supplementary figure S4

#### Slide 1
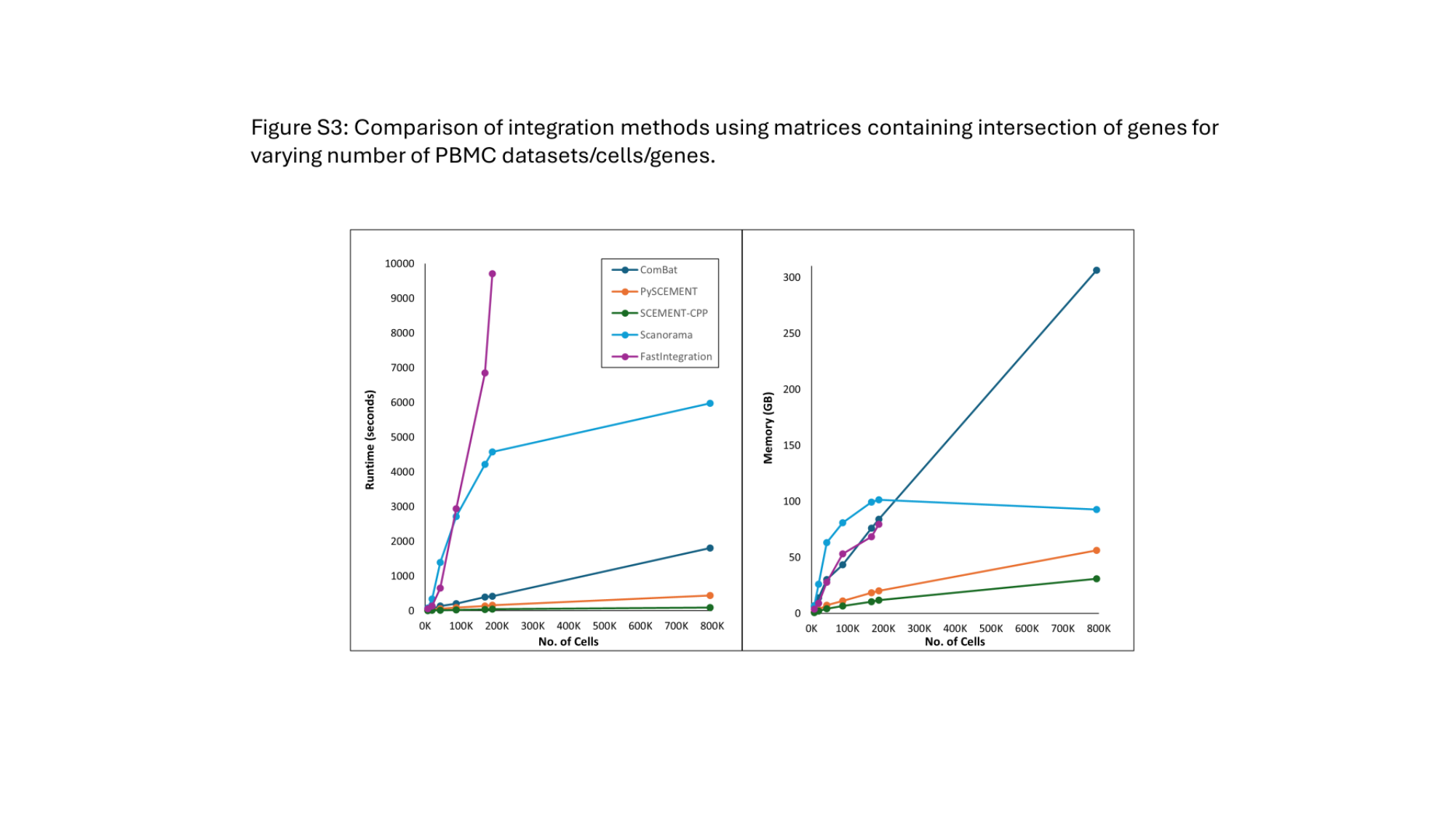

### Supplementary figure S5

#### Slide 1
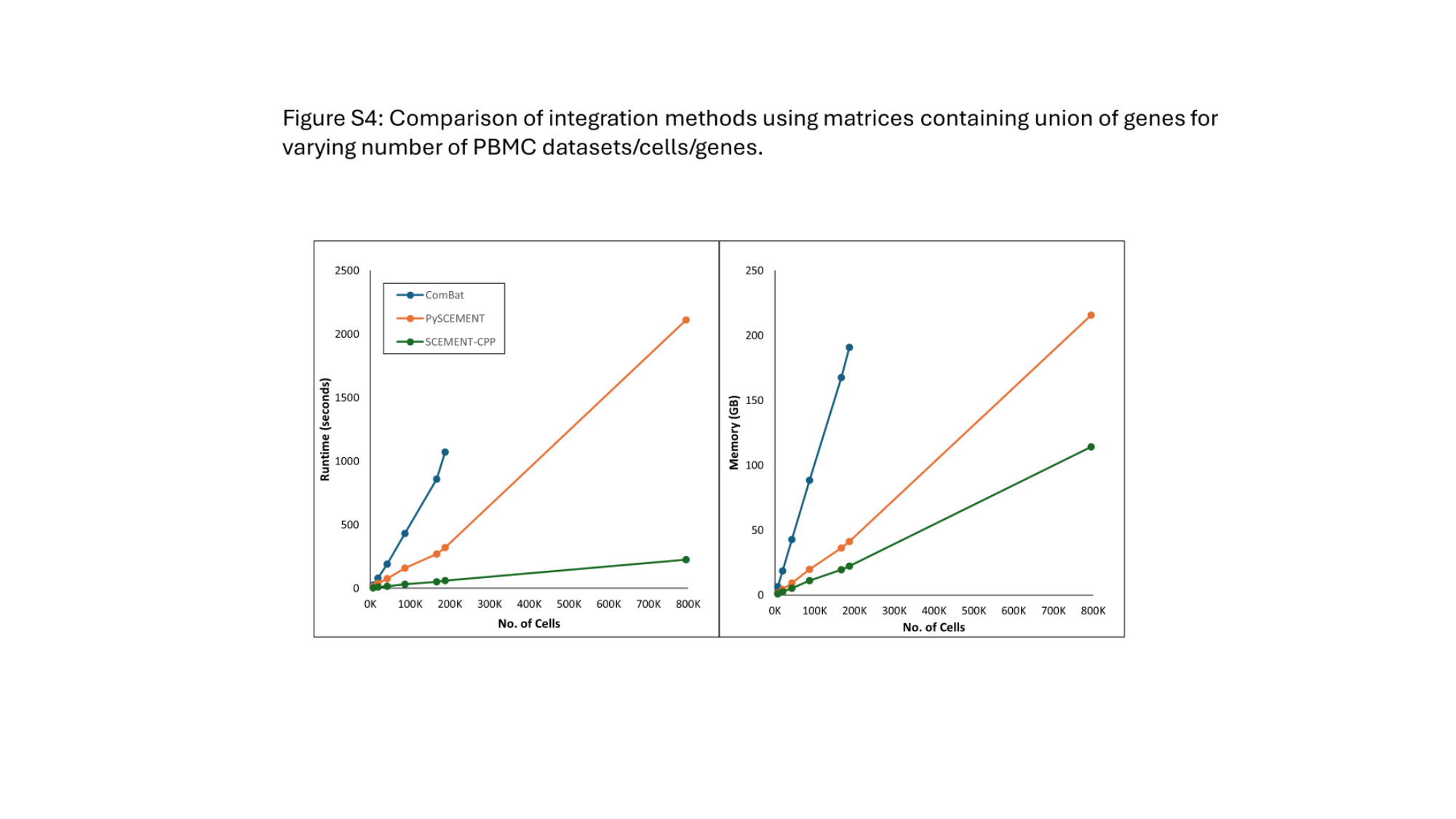
